# Tumour-driven lipid accumulation in oenocytes reflects systemic lipid alterations

**DOI:** 10.1101/2025.07.01.662522

**Authors:** Chang Liu, Sofya Golenkina, Louise Y Cheng

**Affiliations:** Peter MacCallum Cancer Centre, Melbourne, VIC 3000, Australia; Sir Peter MacCallum Department of Oncology, The University of Melbourne, VIC 3010, Australia; School of Medicine, Tsinghua Medicine, Tsinghua University, Beijing, China; Department of Anatomy and Physiology, The University of Melbourne, VIC 3010, Australia

**Keywords:** *Drosophila*, oenocytes, cancer cachexia

## Abstract

Cancer cachexia is a multifactorial syndrome characterized by systemic metabolic dysfunction, including liver steatosis. In this study, we examined the role of larval oenocytes - hepatocyte-like cells, in a *Drosophila* model of cancer cachexia. We found that oenocytes in tumour-bearing larvae accumulate lipid droplets in response to tumour-secreted signals, Gbb and ImpL2. This lipid accumulation reflects systemic changes in lipid metabolism, responding to lipid metabolism manipulations in either the fat body or the muscle. Disrupting lipid synthesis (via FASN1 and DGAT1), storage (via Lsd2), or trafficking (via apolipoproteins) in these tissues significantly modulated lipid droplet accumulation in oenocytes. Moreover, oenocyte-specific knockdown of FASN1 reduced their lipid content and non-autonomously affected lipid droplet size in the fat body, suggesting cross-regulatory interactions between these tissues. Cachectic oenocytes also exhibited altered signaling profiles, characterized by reduced PI3K and elevated Wnt and Ecdysone activity. Enhancing PI3K signaling through Akt overexpression restored oenocyte size and reduced lipid levels; however, these changes did not significantly improve muscle integrity. Together, our data suggests that dynamic exchange of lipids occur between the fat body, oenocytes and the muscle during cancer cachexia. While the fat body and muscle lipid pools are key regulators of muscle integrity, oenocytes - despite their metabolic responsiveness, do not appear to play an active role in preserving muscle function during cachexia.

## Introduction

Cachexia is a multi-factorial, heterogeneous wasting disease affecting around 30% of all cancer patients and around 80% of advanced cancer patients (Baracos *et al*, 2018; Fearon *et al*, 2012). There is so far no gold standard for cachexia treatment, due to a lack of basic mechanistic understanding of the disease. The most prominent manifestation of the disease involves muscle and fat wasting (Baracos *et al*, 2018; Argilés *et al*, 2014), however, it is known that other metabolically active organs such as the bones, brain, liver, gut and heart are likely also involved in this complex inter-organ communication network (Liu *et al*, 2022). Taking advantage of the unparalleled genetic tractability of *Drosophila*, we and others have developed several larval and adult models of cachexia (Bakopoulos *et al*, 2023; Newton *et al*, 2020; Figueroa-Clarevega & Bilder, 2015; Kwon *et al*, 2015; Lee *et al*, 2021; Song *et al*, 2019; Santabárbara-Ruiz & Léopold, 2021; Lodge *et al*, 2021; Khezri *et al*, 2021; Dark *et al*, 2024; Ding *et al*, 2021). These tools allow rapid identification and functional rescue of metabolic and signalling defects using tissue specific drivers.

During cancer cachexia, notable alterations occur in the liver, such that patients can suffer from hepatomegaly and hepatic fibrosis (Rosa-Caldwell *et al*, 2020). These symptoms are attributed to the disruption of hepatic metabolism. Under cachexia, the liver secretes fewer lipids and gluconeogenesis is elevated through the utilisation of amino acid from muscle wasting (Baracos *et al*, 2018). Oenocytes are groups of cells located on both sides of each abdominal segment (Makki *et al*, 2014), together with the fat body, they play equivalent functions as that of the human hepatocytes (Gutierrez *et al*, 2007). While the main function of the fat body is energy storage and utilization, the larval oenocytes are responsible for lipid mobilization, molting and tracheal waterproofing (Parvy *et al*, 2012; Gutierrez *et al*, 2007). Enzymes for lipid-related synthesis and catabolism such as Lipophorin receptors, acetyl-CoA carboxylase (ACC), fatty acid synthase (FAS), and fatty acid β-oxidation enzymes are all enriched in the oenocytes (Makki *et al*, 2014). Therefore, these cells have been implicated in regulating lipid dynamics during nutrient restriction (Gutierrez *et al*, 2007). Furthermore, larval oenocyte-derived hydrocarbons are essential for the waterproofing of the insect trachea (Parvy *et al*, 2012). Oenocyte ablation or the knockdown of Very Long Chain Fatty Acid (VLCFA) enzymes (*ACC, KAR, elongase*), result in severe tracheal defects, with the tracheal tubes filled with aqueous solution (Parvy *et al*, 2012).

As liver steatosis is a key feature of cancer cachexia, in this study, we examined the involvement of larval oenocytes in our *Drosophila* cachexia model. We asked if oenocytes underwent metabolic and signalling alteration during cancer cachexia, and whether these alterations can drive metabolic and/or functional changes in muscle and adipose tissues. We found that oenocytes in tumour-bearing larvae accumulate lipid droplets, a phenotype dependent on tumour-derived signals. Lipid accumulation in the oenocytes reflects altered lipid changes in the fat body. Manipulating lipid metabolism (via *FASN1*, *DGAT1 and Lsd2*) or lipid trafficking (via *Apolipoprotein*) in the fat body or the muscles of tumour bearing animals significantly influenced lipid accumulation in the oenocytes. Furthermore, oenocyte-specific knockdown of *FASN1* reduced lipid accumulation within oenocytes and, non-autonomously, altered lipid droplet size in the fat body. These findings suggest an exchange of lipid pools in the fat body, muscles and the oenocytes. We also found that oenocytes altered signalling in cachectic animals, including reduced PI3K, and upregulated Wnt and Ecdysone signalling. Increasing PI3K signalling via Akt overexpression was able to increase oenocyte size and reduce lipid accumulation, as well as increase fat body size and reduce lipid droplet size in the fat body. However, oenocyte-specific manipulations of lipid metabolism or signalling did not significantly alter muscle integrity, indicating that muscle wasting occurs either upstream or in parallel to the regulatory mechanisms of the oenocytes.

## Results

### Oenocyte lipid droplet accumulation is dependent on tumour secreted factors Gbb and ImpL2

As liver steatosis is a key feature of cancer cachexia, we began by examining the involvement of larval oenocytes in our *Drosophila* cachexia models. We utilise two models of cancer cachexia previously established in the lab (Lodge *et al*, 2021) (Figure 1A) to study the lipid accumulation phenotype. In both models (*Ras^V12^dlg1^RNAi^*and *Ras^V12^scrib^RNAi^*), we observed a robust accumulation of lipid droplets (LDs) in oenocytes, marked by ACC, beginning at day 6 after egg laying (AEL) (Figure 1B–E, G–I; quantified in F and J). Notably, this accumulation occurs one day after lipid droplet enlargement is first detected in the fat body at day 5 AEL (Lodge *et al*, 2021), suggesting that oenocyte lipid accumulation may lie downstream of lipid alterations in the fat body.

**Figure 1.**
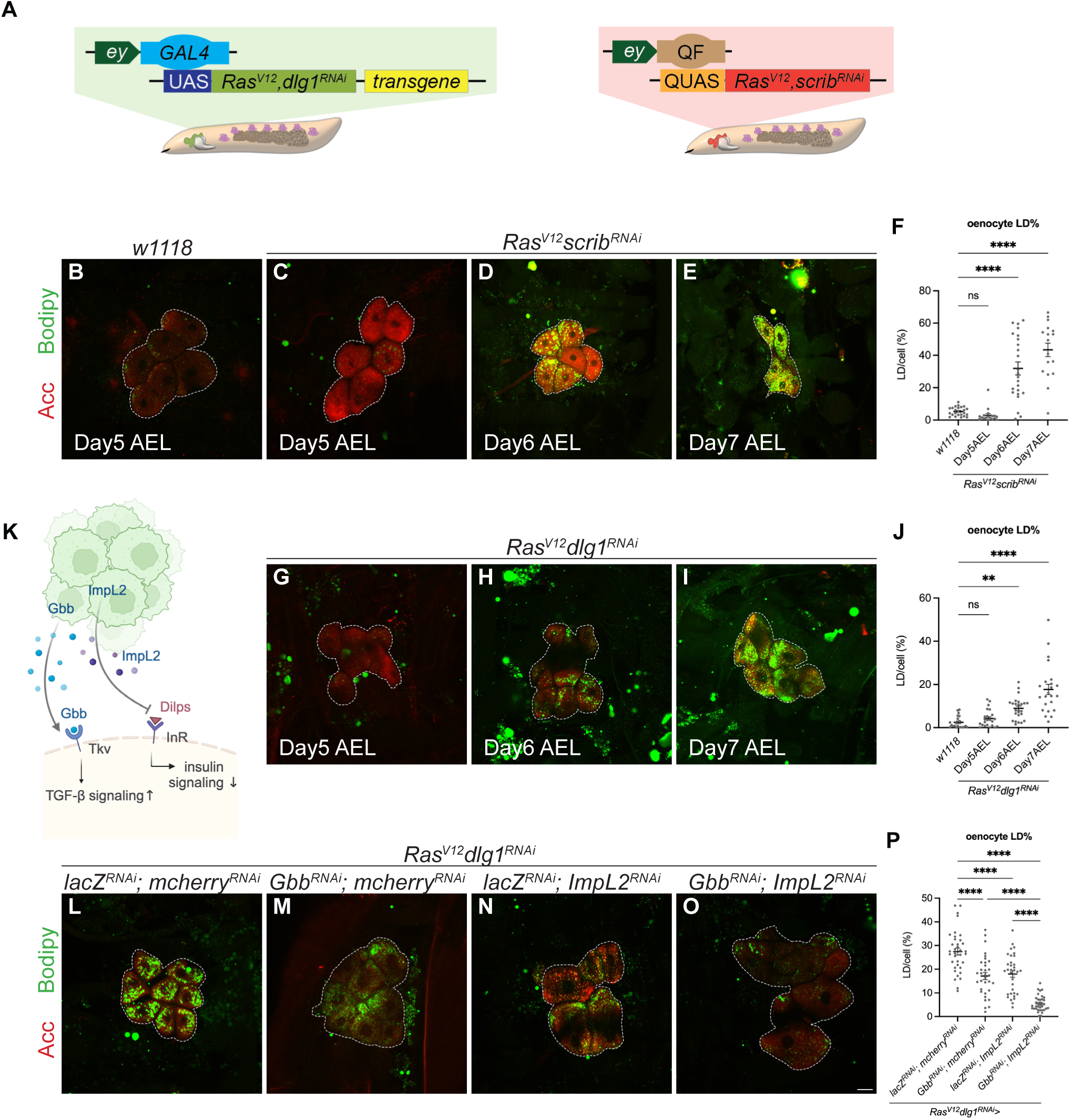
Lipid droplets accumulate in larval oenocytes in response to tumour secreted Impl2 and Gbb. **(A)** Schematics showing *Drosophila* larval tumour models utilised in this study, depicting tumour (green or red), oenocytes (purple) and fat body (light brown). The tumour is induced via *ey-GAL4* (left) or *ey-QF* (right) driving the expression of *UAS-Ras^V12^ dlg1^RNAi^* (left) or *QUAS-Ras^V12^ scrib^RNAi^* (right). The *Ras^V12^ dlg1^RNAi^* model (left) is also used to perform tumour-specific knockdown of transgenes of interest. **(B-E)** Representative maximum projection of the oenocytes (dashed lines) from Day5 AEL *w1118* (B) and Day5-7 AEL *Ras^V12^ scrib^RNAi^* tumour-bearing animals (C-E, respectively). Oenocytes marked by Acc (red), and neutral lipid droplets labelled with Bodipy (green). **(F)** Quantification of LD area as a percentage of individual oenocyte cell area in (B-E). One-way ANOVA: ns p = 0.8941, **** p < 0.0001. Day5 AEL *w1118*: n = 24, mean ± SEM = 5.313 ± 0.5560. Day5 AEL *Ras^V12^ scrib^RNAi^*: n = 18, mean ± SEM = 2.878 ± 1.006. Day6 AEL *Ras^V12^ scrib^RNAi^*: n = 24, mean ± SEM = 31.87 ± 4.011. Day7 AEL *Ras^V12^ scrib^RNAi^*: n = 17, mean ± SEM = 43.42 ± 4.210. **(G-I)** Representative maximum projections of the oenocytes (dashed lines) from Day5-7 AEL *Ras^V12^ dlg1^RNAi^* tumour-bearing animals. Acc (red), Bodipy (green). **(J)** Quantification of LD area as a percentage of individual oenocyte cell area in (G-I). One-way ANOVA: ns p = 0.7146, ** p = 0.0053, **** p < 0.0001. Day5 AEL *w1118*: n = 21, mean ± SEM = 2.407 ± 0.6049. Day5 AEL *Ras^V12^ dlg1^RNAi^*: n = 23, mean ± SEM = 4.177 ± 0.8188. Day6 AEL *Ras^V12^ dlg1^RNAi^*: n = 24, mean ± SEM = 8.908 ± 1.064. Day7 AEL *Ras^V12^ dlg1^RNAi^*: n = 23, mean ± SEM = 17.68 ± 2.393. **(K)** Schematic depicting the mechanism of tumour-secreted factors ImpL2 and Gbb affecting cancer cachexia. **(L-O)** Representative maximum projections of the oenocytes (dashed lines) from *Ras^V12^ dlg1^RNAi^* tumour-bearing animals, where *lacZ^RNAi^; mcherry^RNAi^* (L), *Gbb^RNAi^; mcherry^RNAi^* (M), *lacZ^RNAi^; ImpL2^RNAi^* (N), *Gbb^RNAi^; ImpL2^RNAi^* (O) were expressed in the tumour. Acc (red), Bodipy (green). **(P)** Quantification of LD area as a percentage of individual oenocyte cell area in (L-O). One-way ANOVA: **** p < 0.0001. *lacZ^RNAi^; mcherry^RNAi^*: n = 36, mean ± SEM = 27.46 ± 1.482. *Gbb^RNAi^; mcherry^RNAi^*: n = 33, mean ± SEM = 17.14 ± 1.482. *lacZ^RNAi^; ImpL2^RNAi^*: n = 34, mean ± SEM = 18.00 ± 1.402. *Gbb^RNAi^; ImpL2^RNAi^*: n = 36, mean ± SEM = 5.509 ± 0.5403. Scale bar is 25μm for oenocytes.

As tumour-secreted factors are the drivers of most of the visible disruptions in cachexia (Lodge *et al*, 2021), we next assessed if tumour-secreted factors were responsible for the oenocyte lipid accumulation phenotype. To do so, we specifically knocked down the previously identified tumour secreted factors TGF-beta ligand Gbb and IGF binding protein ImpL2 in the tumour (Figure 1A, K). The knockdown of either factor was able to significantly rescue the lipid accumulation phenotype in the oenocytes (Figure 1L-N, quantified in P), and the knockdown of both factors further reduced LD accumulation in the oenocytes (Figure 1O, quantified in P). These results reinforced the idea that these two tumour-secreted factors function in parallel to affect oenocyte lipid accumulation during cancer cachexia.

### Lipid accumulation in the oenocytes occurs downstream of fat body lipid synthesis/storage/transport

Oenocytes have been shown to play a role in lipid storage upon nutrient restriction, and animals with oenocyte ablation fail to survive during nutrient restriction (Gutierrez *et al*, 2007). Next, we tested if altering lipid synthesis, or lipid storage in the fat body affects lipid accumulation in the oenocytes. Fatty acid synthetase 1 is an enzyme that catalyzes the de novo biosynthesis of fatty acids from acetyl-CoA (Figure 2B). To assess the impact of fat body lipid metabolism on oenocyte lipid accumulation, we specifically knocked down FASN1 in the fat body of tumour-bearing (*Ras^V12^scrib^RNAi^*) larvae (Figure 2A). This led to a significant reduction in lipid droplet size within the fat body (Figure 2C, F) and a marked decrease in lipid accumulation in the oenocytes (Figure 2D, G; quantified in M). Notably, this manipulation also improved muscle integrity (Figure 2E, H; quantified in N). Similarly, fat body-specific knockdown of DGAT1, which catalyses the conversion of diacylglycerol (DAG) to triacylglycerol (TAG) (Cases *et al*, 1998) (Figure 2B), also significantly reduced lipid droplet size in the fat body (Figure 2I, K) and led to a corresponding reduction in oenocyte lipid accumulation (Figure 2J, L; quantified in O). Together, these results indicate that disrupting lipid synthesis and/or storage in the fat body is sufficient to suppress oenocyte lipid accumulation in cachectic animals.

**Figure 2.**
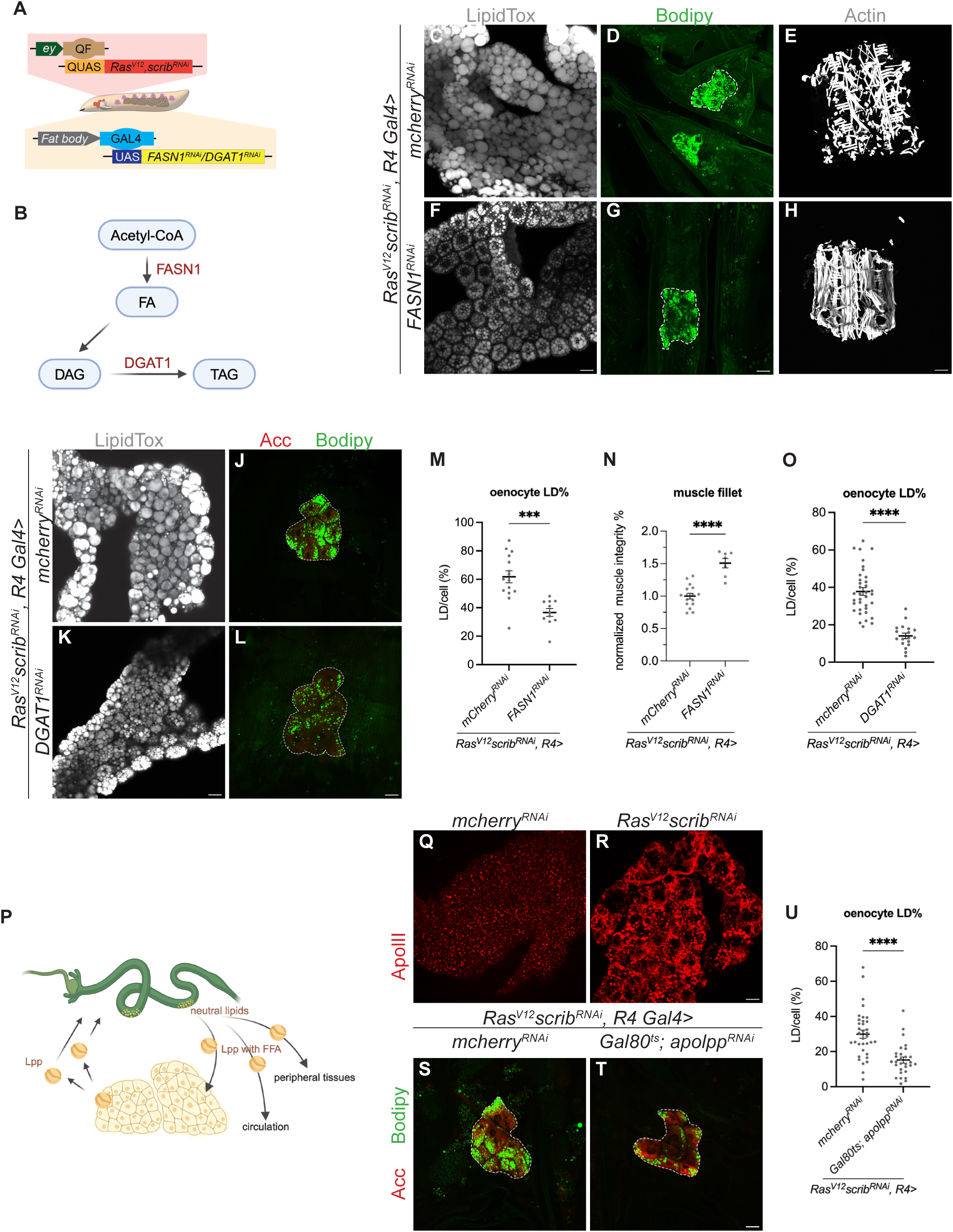
LD accumulation in the oenocytes occurs downstream of fat body lipid synthesis/storage/transport. **(A)** Schematic depicting the dual expression system utilised in this study. *QF-QUAS* induced *Ras^V12^ scrib^RNAi^* tumour in imaginal eye disc while *GAL4-UAS* drove transgene of interest in the fat body. **(B)** Schematic depicting the simplified TAG synthesis pathway. **(C, F)** Representative maximum projections of the LDs in fat body from *Ras^V12^ scrib^RNAi^* tumour-bearing animals, where *mcherry^RNAi^* (C), *FASN1^RNAi^* (F) were expressed in the fat body. LDs labelled by LipidTox (grey). **(D, G)** Representative maximum projections of the oenocytes (dashed lines) from *Ras^V12^ scrib^RNAi^* tumour-bearing animals, where *mcherry^RNAi^* (J), *FASN1^RNAi^* (L) were expressed in the fat body. Bodipy (green). **(E, H)** Representative images of the muscle fillet from *Ras^V12^ scrib^RNAi^* tumour-bearing animals, where *mcherry^RNAi^* (E), *FASN1^RNAi^* (H) were expressed in the fat body. Muscles are visualised by the Phalloidin staining of Actin filaments (grey). **(I, K)** Representative maximum projections of the LDs in fat body from *Ras^V12^ scrib^RNAi^* tumour-bearing animals, where *mcherry^RNAi^*(I), *DGAT1^RNAi^* (K) were expressed in the fat body. LDs labelled by LipidTox (grey). **(J, L)** Representative maximum projections of the oenocytes (dashed lines) from *Ras^V12^ scrib^RNAi^* tumour-bearing animals, where *mcherry^RNAi^*(J), *DGAT1^RNAi^* (L) were expressed in the fat body. Acc (red), Bodipy (green). **(M)** Quantification of LD area as a percentage of individual oenocyte cell area in (D, G). Two-tailed unpaired t test: *** p = 0.0001. *mcherry^RNAi^*: n = 15, mean ± SEM = 61.78 ± 4.155. *FASN1^RNAi^*: n = 11, mean ± SEM = 36.65 ± 2.791. **(N)** Quantification of normalized muscle detachment in (E, H). Two-tailed unpaired t test: **** p < 0.0001. *mcherry^RNAi^*: n = 16, mean ± SEM = 1.000 ± 0.04289. *FASN1^RNAi^*: n = 7, mean ± SEM = 1.507± 0.07295. **(O)** Quantification of LD area as a percentage of individual oenocyte cell area in (J, L). Two-tailed unpaired t test: **** p < 0.0001. *mcherry^RNAi^*: n = 36, mean ± SEM = 37.76 ± 2.046. *DGAT1^RNAi^*: n = 18, mean ± SEM = 14.12 ± 1.480. **(P)** Schematic depicting the production of Lipoprotein from fat body and the lipid transporting process via Lpp. **(Q-R)** Representative maximum projections of the fat body from *mcherry^RNAi^* (B) and *Ras^V12^ scrib^RNAi^* tumour-bearing animals (C). Fat body stained for ApolII (red). **(S, T)** Representative maximum projections of the oenocytes (dashed lines) from *Ras^V12^ scrib^RNAi^* tumour-bearing animals raised at 18°C for 5 days following 29°C for 3 days, where *mcherry^RNAi^* (I), *Gal80^ts^;apolpp^RNAi^*(M) were expressed in the fat body. Acc (red), Bodipy (green). **(U)** Quantification of LD area as a percentage of individual oenocyte cell area in (S, T). Mann-Whitney test: **** p < 0.0001. *mcherry^RNAi^*: n = 35, mean ± SEM = 29.90 ± 2.312. *Gal80^ts^;apolpp^RNAi^*: n = 29, mean ± SEM = 15.18 ± 1.688. Scale bar is 25μm for oenocytes and fat body, 250μm for muscle fillet.

Next, we tested whether inter-organ lipid trafficking plays a role in oenocyte lipid accumulation. Lipophorins (Lpp) are the major lipoproteins in flies (Palm *et al*, 2012), they are synthesized and secreted by fat body cells, and the loss of fat body Lpp production or secretion perturbs inter-organ nutrient flux, causing lipid accumulation in the mid-gut and developmental arrest (Palm *et al*, 2012; Matsuo *et al*, 2019) (Figure 2P). We hypothesise that disruptions in apolipophorins (Apolpp), the scaffolding proteins in the Lipophorin complex, may contribute to the ectopic lipid accumulation phenotype in the oenocytes (Palm *et al*, 2012). As Apolpp is exclusively synthesized by the fat body, we first examined its expression in wild-type and tumour-bearing animals. We observed a significant upregulation of Apolpp in the fat body of cachectic larvae (Figure 2Q, R). To assess whether this elevation contributes to lipid droplet (LD) accumulation in oenocytes, we knocked down apolpp specifically in the fat body of tumour-bearing animals using the fat body-specific driver R4-Gal4 (Supplementary Figure 1A, B). The expression of *apolpp^RNAi^* in the fat body from the beginning of development caused early larval lethality. To bypass this, we temporally expressed *apolpp^RNAi^* for 5 days from L2 using the *GAL80^ts^* system and found this manipulation significantly reduced lipid accumulation in the oenocytes (Figure 2S, T, quantified in U), suggesting that the inhibition of lipid transport from the fat body prevents lipid accumulation in the oenocytes during cancer cachexia.

### Muscle-specific alterations in lipid synthesis/storage caused changes in LD accumulation in the oenocytes

Together, our data suggest that lipid accumulation in oenocytes occurs downstream of lipid synthesis (FASN1), storage (DGAT1), and transport (Apolpp) in the fat body. To further explore inter-tissue lipid communication, we next asked whether modulating lipid stores in the muscle could similarly influence oenocyte lipid accumulation. Using MHC-GAL4 (Figure 3A, (Dark *et al*, 2024)), we inhibited lipid synthesis by knocking down FASN1 specifically in the muscles of the tumour bearing animals. This manipulation did not significantly alter the average LD area in the fat body (Figure 3 B, C, quantified in H), but did result in a drastic reduction in lipid accumulation in the oenocytes (Figure 3 E, F, quantified in I). To test this, we overexpressed Lsd-2, the Drosophila homolog of perilipin 2 in the muscles (Bi *et al*, 2012). This manipulation led to a significant increase in lipid droplet size in the fat body (Figure 3B, D; quantified in H), accompanied by a marked increase in lipid accumulation in the oenocytes (Figure 3E, G; quantified in I).

**Figure 3.**
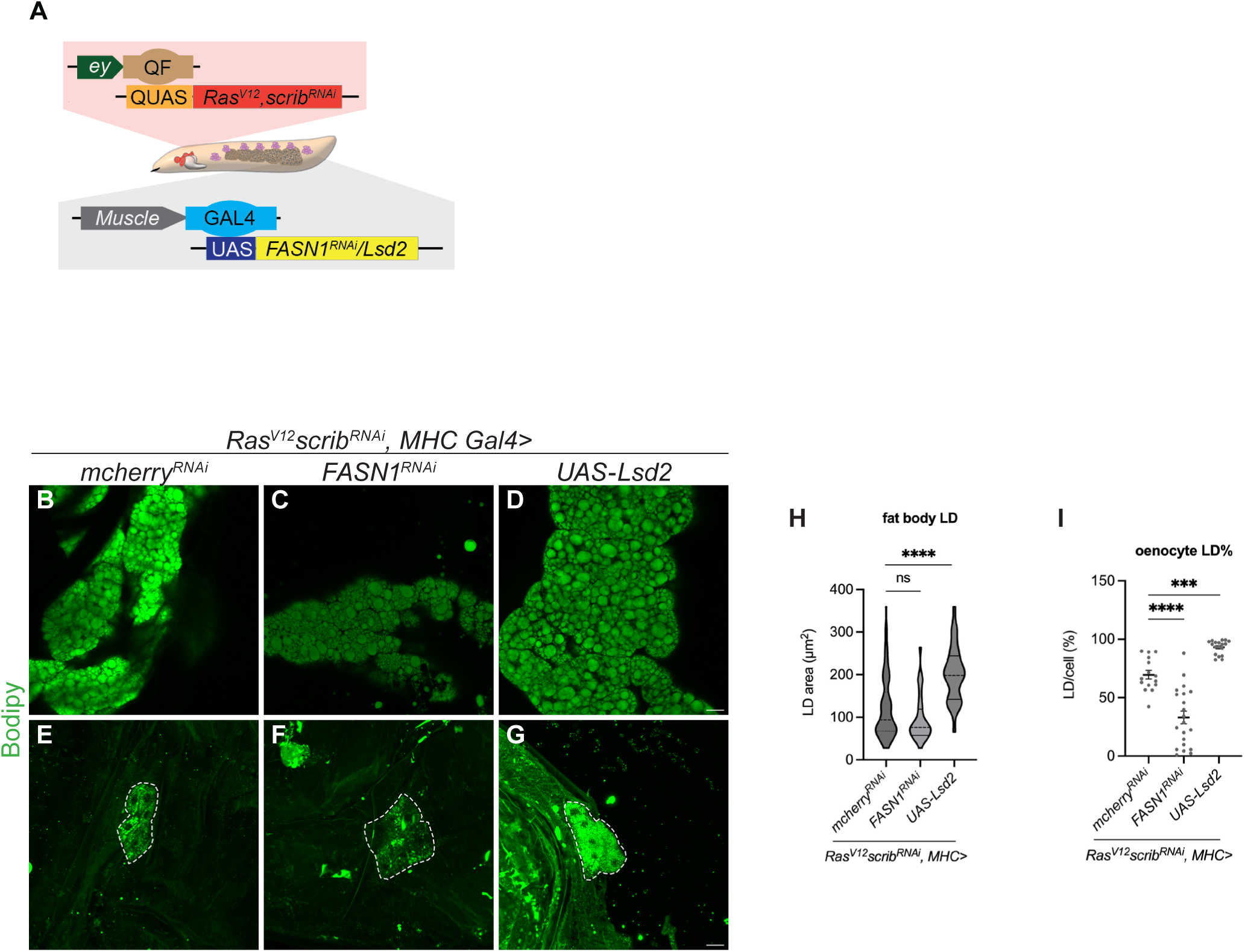
Muscle-specific alterations in lipid synthesis/storage caused changes in LD accumulation in the oenocytes. **(A)** Schematic depicting the dual expression system utilised in this study. *QF-QUAS* induced *Ras^V12^ scrib^RNAi^* tumour in imaginal eye disc while *GAL4-UAS* drove transgene of interest in the muscle. **(B-D)** Representative maximum projections of the LDs in fat body from *Ras^V12^ scrib^RNAi^* tumour-bearing animals, where *mcherry^RNAi^*(B), *FASN1^RNAi^* (C), *UAS-Lsd2* (D) were expressed in the muscle. LDs labelled by Bodipy (green). **(E-G)** Representative maximum projections of the oenocytes (dashed lines) from *Ras^V12^ scrib^RNAi^* tumour-bearing animals, where *mcherry^RNAi^*(E), *FASN1^RNAi^* (F), *UAS-Lsd2* (G) were expressed in the muscle. Bodipy (green). **(H)** Quantification of LD area in fat body in (B-D). Kruskal-Wallis test and Dunn’s test to correct for multiple comparisons: ns p = 0.1467, **** p < 0.0001. *mcherry^RNAi^*: n = 125, mean ± SEM = 120.0 ± 6.332. *FASN1^RNAi^*: n = 36, mean ± SEM = 95.82 ± 8.856. *UAS-Lsd2*: n = 33, mean ± SEM = 201.9 ± 11.25. **(I)** Quantification of LD area as a percentage of individual oenocyte cell area in (E-G). One-way ANOVA: *** p = 0.0007, **** p < 0.0001. *mcherry^RNAi^*: n = 15, mean ± SEM = 69.58 ± 3.670. *FASN1^RNAi^*: n = 21, mean ± SEM = 33.01 ± 5.497. *UAS-Lsd2*: n = 18, mean ± SEM = 93.02 ± 1.400. Scale bar is 25μm for oenocytes and fat body.

### Oenocyte-specific inhibition of lipid synthesis reduces LD accumulation in the oenocytes and the fat body

Next, we investigated whether disrupting lipid synthesis directly within oenocytes could modulate their lipid accumulation and, in turn, influence fat body and muscle homeostasis in cachectic animals. Using *promE-Gal4* (specifically expressed in the oenocytes, Supplement Figure 1C, D, quantified in E), we knocked down *FASN1* (Figure 2B) specifically in the oenocytes of tumour-bearing animals (Figure 4A). Oenocyte-specific knockdown of FASN1 led to a significant reduction in lipid droplets within oenocytes (Figure 4B, E; quantified in H). Interestingly, this manipulation also decreased lipid droplet size in the fat body of tumour-bearing animals, but not in controls (Figure 4D, G; quantified in J and Supplementary Figure1F, G), indicating a non-autonomous effect. However, unlike fat body-specific FASN1 knockdown (Figure 2), inhibiting lipid synthesis in oenocytes was not sufficient to improve muscle morphology (Figure 4C, F; quantified in I).

**Figure 4.**
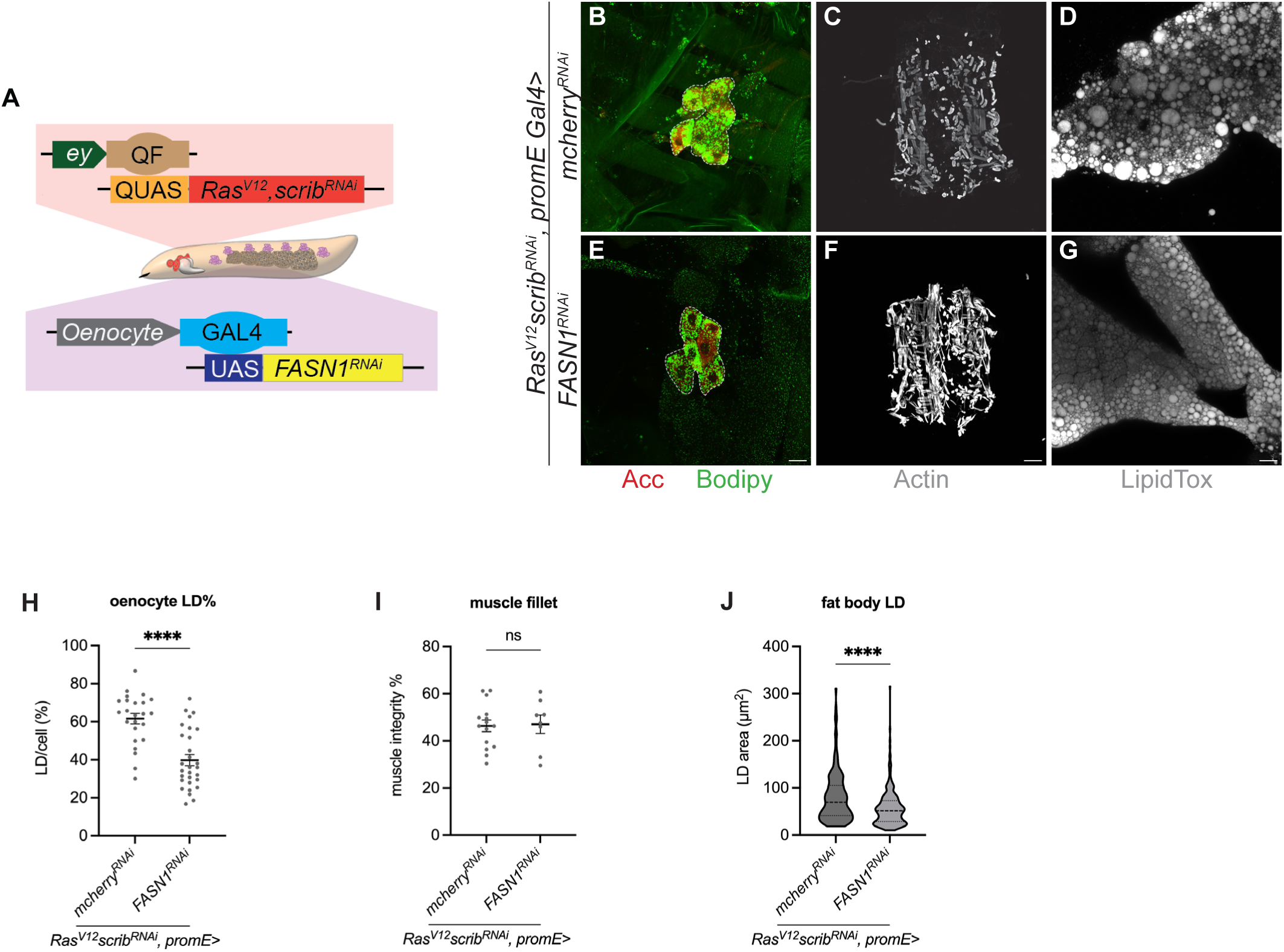
Oenocyte-specific inhibition of lipid synthesis reduces LD accumulation in the oenocytes and the fat body. **(A)** Schematic depicting the dual expression system utilised in this study. *QF-QUAS* induced *Ras^V12^ scrib^RNAi^* tumour in imaginal eye disc while *GAL4-UAS* drove transgene of interest in the oenocytes. **(B, E)** Representative maximum projections of the oenocytes (dashed lines) from *Ras^V12^ scrib^RNAi^* tumour-bearing animals, where *mcherry^RNAi^*(B), *FASN1^RNAi^* (E) were expressed in the oenocytes. Acc (red), Bodipy (green). **(C, F)** Representative images of the muscle fillet from *Ras^V12^ scrib^RNAi^* tumour-bearing animals, where *mcherry^RNAi^* (C), *FASN1^RNAi^* (F) were expressed in the oenocytes. Actin (grey). **(D, G)** Representative maximum projections of the LDs in fat body from *Ras^V12^ scrib^RNAi^* tumour-bearing animals, where *mcherry^RNAi^* (D), *FASN1^RNAi^*(G) were expressed in the oenocytes. LipidTox (grey). **(H)** Quantification of LD area as a percentage of individual oenocyte cell area in (B, E). Two-tailed unpaired t test: **** p < 0.0001. *mcherry^RNAi^*: n = 24, mean ± SEM = 61.51 ± 2.753. *FASN1^RNAi^*: n = 29, mean ± SEM = 39.80 ± 2.889. **(I)** Quantification of muscle detachment in (C, F). Two-tailed unpaired t test: ns p = 0.8831. *mcherry^RNAi^*: n = 15, mean ± SEM = 46.371 ± 2.490. *FASN1^RNAi^*: n = 8, mean ± SEM = 47.03 ± 3.840. **(J)** Quantification of LD area in fat body in (D, G). Mann-Whitney test: **** p < 0.0001. *mcherry^RNAi^*: n = 138, mean ± SEM = 80.25 ± 4.593. *FASN1^RNAi^*: n = 243, mean ± SEM = 57.18 ± 2.511. Scale bar is 25μm for oenocytes and fat body, 250μm for muscle fillet.

Together, these results suggest that while modulating lipid metabolism in the fat body can influence both oenocyte lipid accumulation and muscle integrity, altering lipid synthesis in oenocytes primarily affects lipid dynamics within oenocytes and the fat body, without impacting muscle health in cachectic animals.

### Oenocyte-specific activation of PI3K signaling prevents lipid accumulation in cachectic animals

Lipid accumulation in oenocytes is positively regulated by the putative lipid dehydrogenase Spidey/Kar and negatively regulated by PI3K signaling (Gutierrez *et al*, 2007; Cinnamon *et al*, 2016). Notably, knockdown of target of rapamycin (TOR) or the amino acid transporter slimfast (slif) in the larval fat body enhances lipid accumulation in oenocytes, consistent with a systemic reduction in PI3K pathway activity. To assess whether PI3K signaling is altered in the oenocytes of cachectic animals, we examined the localization of FOXO-GFP, a reporter for PI3K/TOR pathway activity (Dong *et al*, 2025) (Figure 5A). We found that FOXO expression level is significantly increased in the nucleus of oenocytes (normalised to DAPI), indicating a downregulation of the PI3K/Tor signalling pathway (Figure 5B-C’, quantified in D).

**Figure 5.**
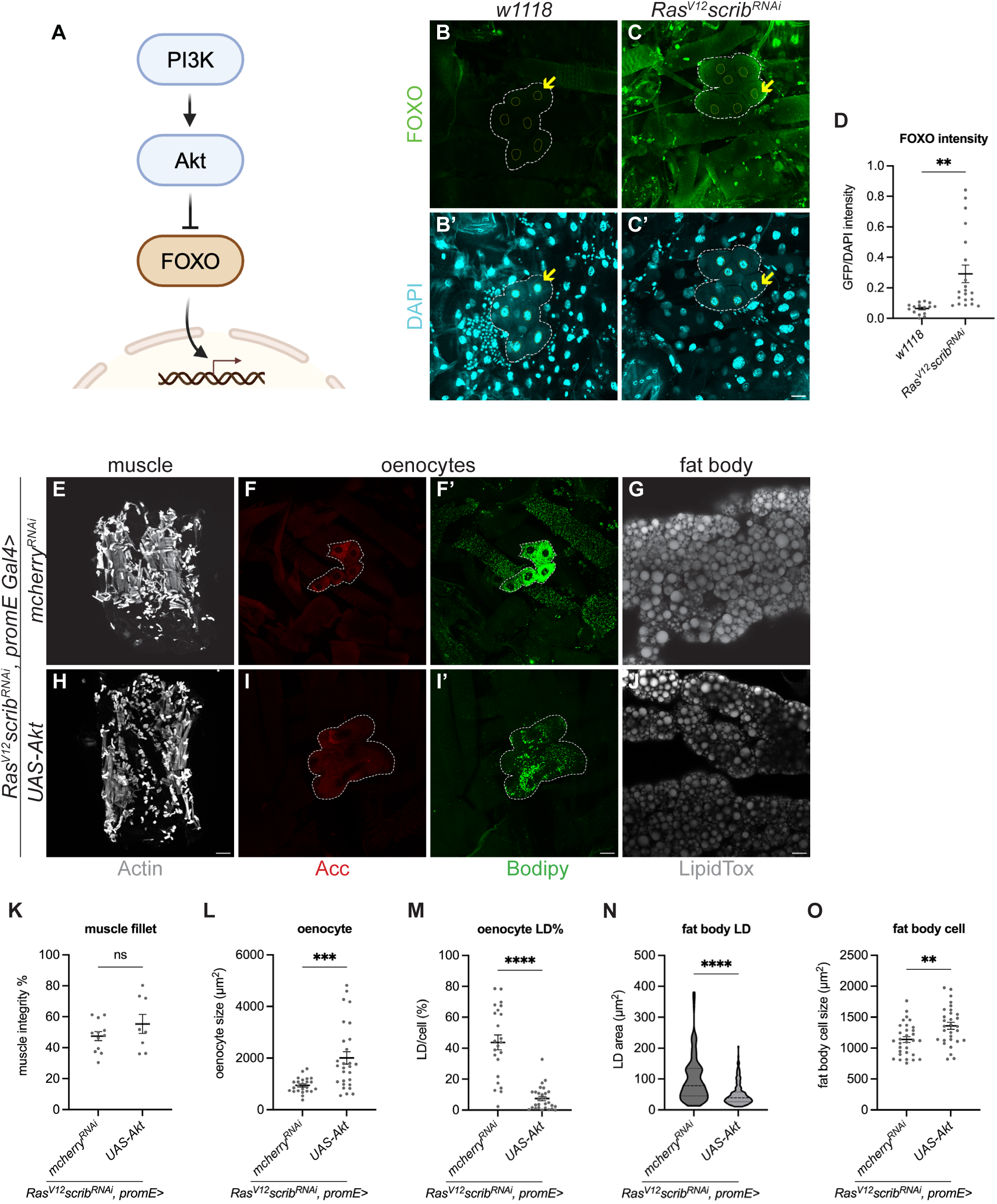
Oenocyte-specific activation of PI3K signaling prevents lipid accumulation in cachectic animals. **(A)** Schematic depicting the PI3K signalling pathway. **(B-C’)** Representative maximum projections of the oenocytes (dashed lines) from *w1118* (B-B’) and *Ras^V12^ scrib^RNAi^* tumour-bearing animals (C-C’). Oenocytes expressed endogenous Foxo-GFP (green) in the nucleus (yellow arrows) (B-C), counterstained with DAPI (cyan) (B’-C’). **(D)** Quantification of oenocyte FoxO-GFP intensity normalised to DAPI intensity in (B-C’). Two-tailed unpaired t test: ** p = 0.0019. *w1118*: n = 15, mean ± SEM = 0.06460 ± 0.009098. *Ras^V12^ scrib^RNAi^*: n = 20, mean ± SEM = 0.2911 ± 0.05738. **(E, H)** Representative images of the muscle fillet from *Ras^V12^ scrib^RNAi^* tumour-bearing animals, where *mcherry^RNAi^* (E), *UAS-Akt* (H) were expressed in the oenocytes. Actin (grey). **(F-F’, I-I’)** Representative maximum projections of the oenocytes (dashed lines) from *Ras^V12^ scrib^RNAi^* tumour-bearing animals, where *mcherry^RNAi^*(F-F’), *UAS-Akt* (I-I’) were expressed in oenocytes. Acc (red), Bodipy (green). **(G, J)** Representative maximum projections of the LDs in fat body from *Ras^V12^ scrib^RNAi^* tumour-bearing animals, where *mcherry^RNAi^*(G), *UAS-Akt* (J) were expressed in the oenocytes. LipidTox (grey). **(K)** Quantification of muscle detachment in (E, H). Two-tailed unpaired t test: ns p = 0.2184. *mcherry^RNAi^*: n = 12, mean ± SEM = 47.54 ± 2.922. *UAS-Akt*: n = 8, mean ± SEM = 55.39 ± 6.192. **(L)** Quantification of oenocyte cell size in (F, I). Two-tailed unpaired t test: *** p = 0.0002. *mcherry^RNAi^*: n = 24, mean ± SEM = 943.3 ± 58.94. *UAS-Akt*: n = 28, mean ± SEM = 2009 ± 240.5. **(M)** Quantification of LD area as a percentage of individual oenocyte cell area in (F’, I’). Mann-Whitney test: **** p < 0.0001. *mcherry^RNAi^*: n = 23, mean ± SEM = 43.74 ± 4.767. *UAS-Akt*: n = 29, mean ± SEM = 7.435 ± 1.387. **(N)** Quantification of LD area in fat body in (G, J). Mann-Whitney test: **** p < 0.0001. *mcherry^RNAi^*: n = 188, mean ± SEM = 97.92 ± 5.277, *UAS-Akt*: n = 210, mean ± SEM = 50.37 ± 2.411. **(O)** Quantification of fat body cell size in (G, J). Two-tailed unpaired t test: ** p = 0.0036. *mcherry^RNAi^*: n = 31, mean ± SEM = 1138 ± 47.05, *UAS-Akt*: n = 30, mean ± SEM = 1360 ± 56.57. Scale bar is 25μm for oenocytes and fat body, 250μm for muscle fillet.

Next, we asked whether increasing PI3K signaling in oenocytes is sufficient to suppress lipid droplet accumulation. Activation of the PI3K pathway via Akt overexpression in oenocytes led to a significant increase in oenocyte size in both wild-type and cachectic animals (Figure 5F, I; quantified in L; Supplementary Figure 1H–I; quantified in L), and a marked reduction in oenocyte lipid accumulation in cachectic animals (Figure 5F′, I′; quantified in M). However, unlike fat body-specific Akt overexpression—which has been shown to rescue muscle integrity (Bakopoulos *et al*, 2023) Akt expression in oenocytes did not improve muscle morphology in cachectic animals (Figure 5E, H; quantified in K).

Interestingly, Akt overexpression in oenocytes not only altered oenocyte lipid content but also non-autonomously reduced LD size in the fat body of tumour-bearing animals (Figure 5G, J; quantified in N). Additionally, it led to an increase in fat body cell size (Figure 5O), an effect that was also observed in wild-type animals (Supplementary Figure 1J–K; quantified in M), suggesting a broader role for oenocyte-derived signals in regulating fat body size.

Together, our data indicate that, similar to other tissues such as muscle and fat body (Lodge *et al*, 2021; Bakopoulos *et al*, 2023; Dark *et al*, 2024), PI3K signaling is downregulated in oenocytes during cachexia. Enhancing PI3K activity in oenocytes is sufficient to increase their size and reduce LD accumulation both locally and in the fat body. However, these changes are not sufficient to restore muscle integrity, suggesting that oenocyte lipid dynamics are not primary regulators of muscle health in cachectic animals.

### Wnt/wingless and ecdysone signalling are altered in the oenocytes of cachectic animals

We next examined several other signalling pathways in the oenocytes of tumour bearing animals. Utilizing signalling readouts, we assessed whether Hippo, Toll, Imd, Notch, Jnk, Wnt/Wg and Ecdysone signalling pathways are significantly altered in the oenocytes of cachectic animals compared to the wildtype (Table 1, significant results marked in blue and red). We found that Broad-complex (BR-C), a readout of the ecdysone signalling pathway (Karim *et al*, 1993) as well as Nkd-lacZ – a readout of Wnt/Wg pathway were upregulated in the oenocytes of cachectic animals compared to the wildtype control (Supplement Figure 2A-B’, D-E’, quantified in C, F).

**Table 1.** Signalling pathways screening in oenocytes of tumour-bearing animals vs. wildtype animals and their necessity for oenocyte and muscle cachectic phenotypes.

| pathways | readout in oenocytes | tumour bearing vs. control | required for oenocyte cacheixa phenotype | required for muscle cachexia phenotype |
| --- | --- | --- | --- | --- |
| PI3K | *foxo-GFP* | down-regulated | Yes | No |
| Wnt | *nkd-lacZ* | up-regulated | No | No |
| Ecdysone | Broad complex | up-regulated | No | No |
| Hippo | CycE | not changed | No | No |
| Toll | Dorsal | not changed | No | No |
| Imd | Relish | not changed | No | No |
| JNK | MMP1 | not changed | No | No |
| Notch | Notch extracellular domain | not changed | No | No |

A key function of oenocytes is their role in the biosynthesis and metabolism of ecdysone, an insect moulting-related hormone. It has previously been shown that *spidey*, which encodes a steroid dehydrogenase, affects the animal’s ability to produce pheromones (Chiang *et al*, 2016). However, the overexpression of a dominant negative form of ecdysone receptor (EcR^DN^) (Cherbas *et al*, 2003) did not significantly alter LD accumulation, nor affect muscle integrity in cachectic animals.

The activation of canonical Wnt/Wg signaling has been shown to promote lipolysis while concurrently inhibiting lipogenesis and fatty acid β-oxidation in both larval and adult fat body. To assess whether Wnt/Wg is functionally involved in lipid metabolism in oenocytes, we knocked down *pangolin (pan)*, a downstream factor of Wnt/Wg signalling (Zhang *et al*, 2017) in the oenocytes of tumour bearing animals. This manipulation did not significantly affect oenocyte lipid morphology nor muscle integrity in tumour bearing animals (Supplement Figure 2G-L, quantified in M-N). These results suggested that despite the alteration of these signalling pathways in cachectic animals, they were not functionally involved in lipid accumulation in the oenocytes, nor are involved in muscle integrity. However, they could be involved in other physiological functions of the oenocytes, which require further exploration in the future.

## Discussion

In *Drosophila* larval models of cancer cachexia, we observed abnormal lipid droplet accumulation in oenocytes. This phenotype appears to be driven by tumour-secreted factors and can be modulated through (i) manipulation of lipid synthesis and/or storage in the cachectic fat body or muscle, (ii) disruption of lipid trafficking from the fat body via ApoLpp, and (iii) regulation of PI3K signaling within oenocytes. While altering lipid metabolism in the fat body or muscle (Figure 2; (Dark *et al*, 2024)) was sufficient to influence muscle integrity during cachexia, modulating PI3K signaling or lipid metabolism specifically in oenocytes affected fat body lipid content but had no effect on muscle morphology. These findings suggest that lipid accumulation in oenocytes reflects systemic lipid availability and likely occurs downstream of, or in parallel with, the mechanisms governing muscle integrity in cachectic animals (Figure 6).

**Figure 6.**
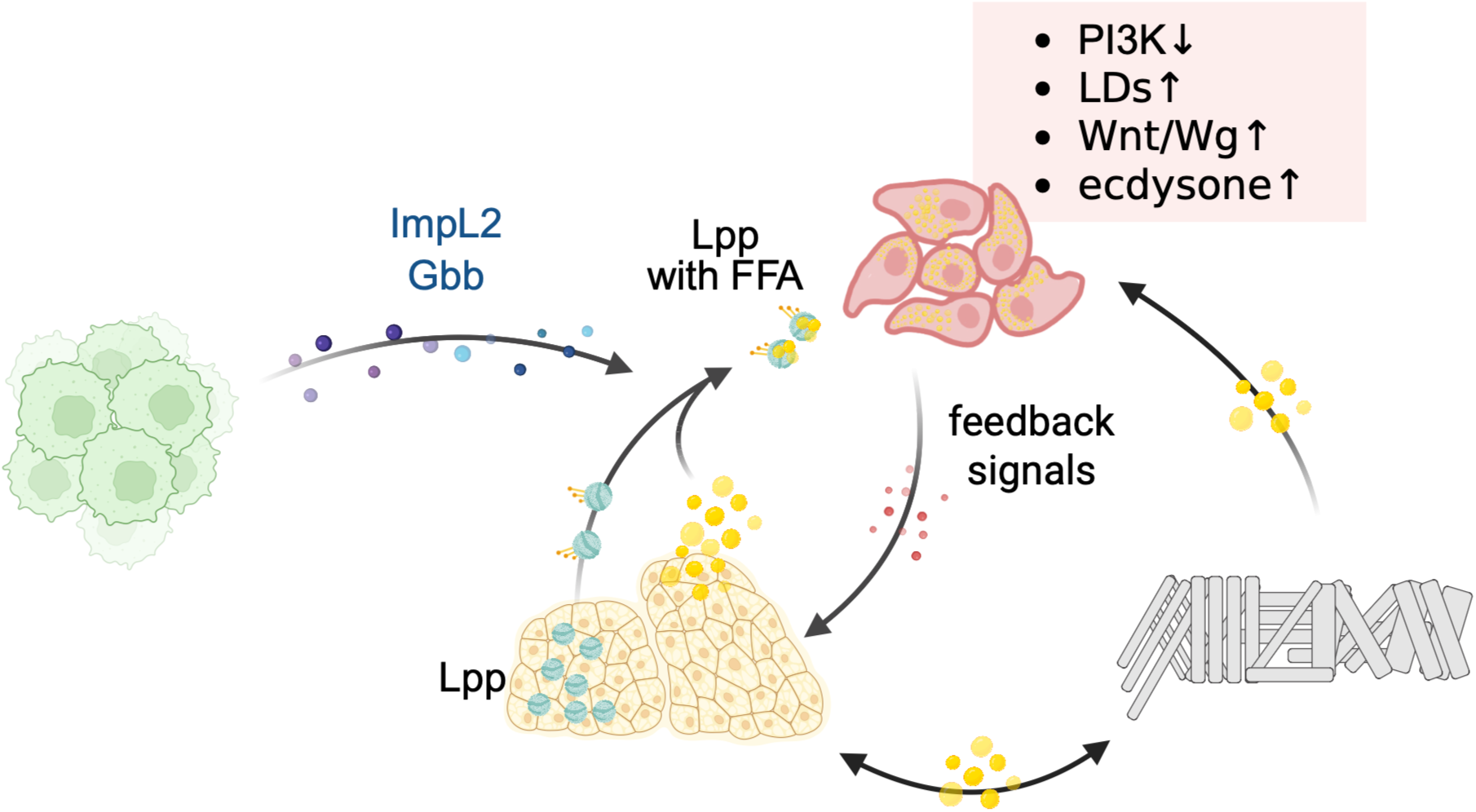
Model of fat body, oenocyte and muscle interaction in cancer cachexia. In cachectic animals, tumours secrete ImpL2 and Gbb, leading to systemic disruption of lipid metabolism. Lipid droplet trafficking occurs dynamically between the fat body, muscle, and oenocytes, with lipid-binding proteins (Lpps) released from the fat body into the hemolymph mediating these lipid exchanges. During cachexia, PI3K signaling in oenocytes is downregulated, whereas Wnt/Wg and ecdysone signaling pathways are upregulated.

Our findings reveal a dynamic exchange of lipids between the fat body, muscles, and oenocytes. We observed that altering lipid synthesis or storage in the fat body and muscle leads to corresponding changes in lipid accumulation within oenocytes. Conversely, manipulating PI3K signaling or lipid synthesis specifically in oenocytes also influenced fat body size and lipid content, as measured by lipid droplet size. Previous work showed that knockdown of Acc in oenocytes can lead to tracheal flooding and hypoxia, which subsequently affects fat body LD morphology (Parvy *et al*, 2012). However, we found no evidence of tracheal flooding in tumour-bearing animals upon FASN1 inhibition in oenocytes (Supplementary Figure 2O, P), suggesting an alternative mechanism. One possibility is that oenocytes can secrete molecules that influence fat body physiology. To investigate this, future studies could employ tissue-specific protein biotinylation (Droujinine *et al*, 2021) to identify candidate proteins that are specifically trafficked between oenocytes and the fat body.

PI3K signaling is deregulated during cancer cachexia, and enhanced PI3K activation significantly reduces ectopic lipid droplet accumulation in cachectic oenocytes. This indicates that PI3K activation is sufficient to suppress LD accumulation in these cells. This observation parallels what occurs in wild-type larvae under nutrient restriction, where PI3K activation similarly abolishes LD accumulation in oenocytes (Cinnamon *et al*, 2016). In adult oenocytes, previous work demonstrated that Pvf1-mediated activation of the PI3K/Akt/TOR pathway suppresses lipid synthesis and reduces LD accumulation (Ghosh *et al*, 2020). However, contrasting data from (Chatterjee *et al*, 2014) showed that activation of insulin/PI3K signaling in adult oenocytes promotes lipid droplet accumulation under both fed and starved conditions. Our findings, in the context of cancer cachexia, align with those of Cinnamon et al. and Ghosh et al. Furthermore, our data suggest that tumour-secreted ligands ImpL2 and Gbb drive oenocyte lipid accumulation. Future studies could investigate whether these ligands induce lipid accumulation through modulation of PI3K or TGF-beta signaling pathways.

In addition to PI3K, we found that Wnt/Wg pathway and Ecdysone-related pathway were upregulated in the cachectic oenocytes. Downregulation of these pathways did not significantly impact on either oenocyte LD accumulation or muscle detachment. In the fat body, severe lipid metabolism defects, including reduced TAG level has been reported to result from hyperactivated Wnt/Wg signalling (Zhang *et al*, 2017). Additionally, the overexpression of *wg* in the muscle led to a depletion of lipid accumulation (Abou Azar & Lim, 2021). Jointly, it remains possible that Wnt/Wg act as a negative regulator in adipogenesis and lipid metabolism in the oenocytes, despite our results. Ecdysone is a key mediator in *Drosophila* for maintaining overall hormonal balance. It has been shown that the circulating ecdysone level was significantly reduced in cachectic animals. During cachexia, the tumour functions as a sink of steroids, contributing to wasting symptoms (Santabárbara-Ruiz & Léopold, 2021). Our results showed that oenocytes exhibited elevated Ecdysone signalling levels as indicated by an ecdysone-inducible reporter - BR-C (Karim *et al*, 1993), it remains possible that a compensatory adaption response happens in the oenocytes. Therefore, in future studies, it will be important to assess whether *br-c* downstream regulators affect oenocytes/fat/muscle phenotypes in the context of cancer cachexia.

## Materials and Methods

### Fly husbandry

The following stocks were used from the Blooming *Drosophila stock Centre*: *UAS-lacZ^RNAi^* (BL31562), *UAS-mcherry^RNAi^* (BL35785), *R4-Gal4* (BL33832), *MHC-Gal4* (BL55133), *promE-Gal4* (on Chr2) (BL65404), *promE-Gal4* (on Chr3) (BL65405), *UAS-Akt* (BL8191), *UAS-pan^RNAi^* (BL26743), *UAS-EcR^DN^* (BL9451), *foxo-GFP* (BL38644), *UAS-apolpp^RNAi^* (BL33388). The following stocks were obtained from the Vienna Drosophila Resource Centre: *UAS-ImpL2^RNAi^* (v30931), *UAS-Gbb^RNAi^* (v330684), *UAS-dgat1^RNAi^* (v100003). The following stocks were also used: *w1118*, *UAS-fasn1^RNAi^* (NIG-Fly #3523R-2), *Ey-FLP1; QUAS-Ras^V12^, QUAS-scrib^RNAi^/CyOQS; act>CD2>QF, UAS-RFP/TMBQS* (Lodge et al, 2021), *Ey-FLP1; UAS-Ras^V12^, UAS-dlg1^RNAi^/CyO, GAL80; act>CD2>GAL4* (Lodge et al, 2021)*, UAS-Lsd2* (Cinnamon et al, 2016).

Fly stocks were reared on standard media. Adults were allowed to lay for 24h at 25°C and the progeny was then moved to 29°C. For experiment with *Ras^V12^ scrib^RNAi^, R4> Gal80^ts^;apolpp^RNAi^*, larvae were reared at 19°C for 5 days and then moved to 29°C for 3 days. Animals were dissected at wandering stage in non-tumour-bearing animals. For tumour-bearing animals, they were dissected as indicated throughout.

### Immunostaining

For muscle fillet staining, larvae were dissected as previously described (Dark *et al*, 2022), fixed for 25 min in PBS containing 4% formaldehyde and washed three times for 5 min each with PBS containing 0.3% Triton-X (PBST-0.3). Fat body samples were fixed for 40 min and washed three times for 5 min each with PBS. Tissues were then stained as per the manufacturer’s specification. Muscle samples were mounted in PBS containing 80% glycerol, fat body samples were mounted in PBS. All samples were imaged on an Olympus FV3000 confocal microscope. Within a given experiment, all images were acquired using identical settings. Primary antibodies used: rabbit anti-Acc (1:50, Cell Signalling #3661), chick anti-β-Galactosidase (1:1000, abcam #ab9361), mouse anti-broad-complex (1:100, DSHB), mouse anti-dorsal (1:10, DSHB), mouse anti-relish (1:20, DSHB), mouse anti-MMP1 (1:100, DSHB), mouse anti-notch-extra (1:50, DSHB), rat anti-CycE (1:200, DSHB), rabbit anti-apolII (1:500, a gift from Dr. Akhila Rajan). Secondary donkey antibodies conjugated to Alexa 488 and Alexa 555 (Molecular Probes) were used at 1:500. Bodipy (Invitrogen), DAPI (Molecular Probes), HCS LipidTOX^TM^ Deep Red Neutral Lipid Stain (Invitrogen #H34477) were used at1:1000. Phalloidin 647 (abcam #ab176759) was used at 1:500.

Muscle fillet phalloidin staining were conducted on day 7 *Ras^V12^scrib^RNAi^* animals and on day 8 *Ras^V12^dlg1^RNAi^*animals. All other fat body and oenocyte staining were conducted on day 6 *Ras^V12^scrib^RNAi^* animals and on day 7 *Ras^V12^dlg1^RNAi^*animals (except when specified in the figure legend).

### Trachea flooding assay

Larvae at the stage 6 hours after L2/L3 molt were transferred from food in vials to PBS with blue food dye in a dish and maintained submerged for 15 minutes (Cinnamon et al, 2016).

### Image analysis

All images were quantified with FIJI. Muscle/cuticle percentage was determined using FIJI as previously described (Dark *et al*, 2022). For muscle atrophy measurements, muscle length and width measurements were done on the VL3 muscle on 20X images of individual fillets with the line selection tool (Bakopoulos et al, 2023). For LD /cell percentage measurement in the oenocyte, a ROI was drawn around the cell outline in the red channel (Acc staining) first. Then the green channel (Bodipy staining) was converted to a binary mask, and the total area of fluorescence detected within the ROI was divided by the total ROI area as % LD/cell. The LD area in the fat body was quantified by drawing a circle around individual lipid droplet within five cells per fat body, followed by area measurement. The foxo-GFP, nkd-lacZ, broad-complex intensity was normalized to DAPI intensity. The fluorescence level was determined using the formula: CTCF (corrected total cell fluorescence) = Integrated density – (Area of selected cell x mean fluorescence of background readings) (McCloy *et al*, 2014).

### Statistical analysis

At least two animals per genotype were used for all experiments. Statistical analyses and graph plotting were all performed using GraphPad Prism. In graphs, error bars represent standard error of mean (SEM). For oenocyte quantifications and cell size quantifications, n = cell number. For muscle integrity or atrophy quantifications, n = animal number. For LD area quantification in fat body, n = LD number. For experiments comparing two conditions, significant differences were tested by two-tailed unpaired t-tests for normally distributed data, or non-parametric Mann-Whitney tests for non-normally distributed data. When comparing more than two conditions, significant differences were tested either by ordinary one-way ANOVA when the data were normally distributed, or by Kruskal-Wallis tests when the data were not normally distributed. Dunnet or Dunn’s tests were used to correct for multiple comparisons following one-way ANOVA and Krustal-Wallis tests, respectively. (ns) p > 0.05, (*) p < 0.05, (**) p < 0.01, (***) p < 0.001, (****) p < 0.0001.

## Acknowledgements

We are grateful to Helena Richardson and Kieran Harvey for the generous sharing of antibodies and fly stocks. We thank the Bloomington *Drosophil*a Stock Centre (BDSC), Vienna *Drosophila* Resource Centre (VDRC), and Developmental Studies Hybridoma Bank (DSHB) for fly stocks and antibodies. We would like to thank OZDros for *Drosophila* quarantine, Peter MacCallum Centre for Advanced Histology and Microscopy for microscopy assistance. Chang Liu is funded by the China Scholarship Council, and was a part of the Tsinghua Medicine-University of Melbourne MRes program. LYC’s laboratory is supported by funding from the NHMRC Ideas Grant (APP2011289).

---

**Supplementary Figure 1.**
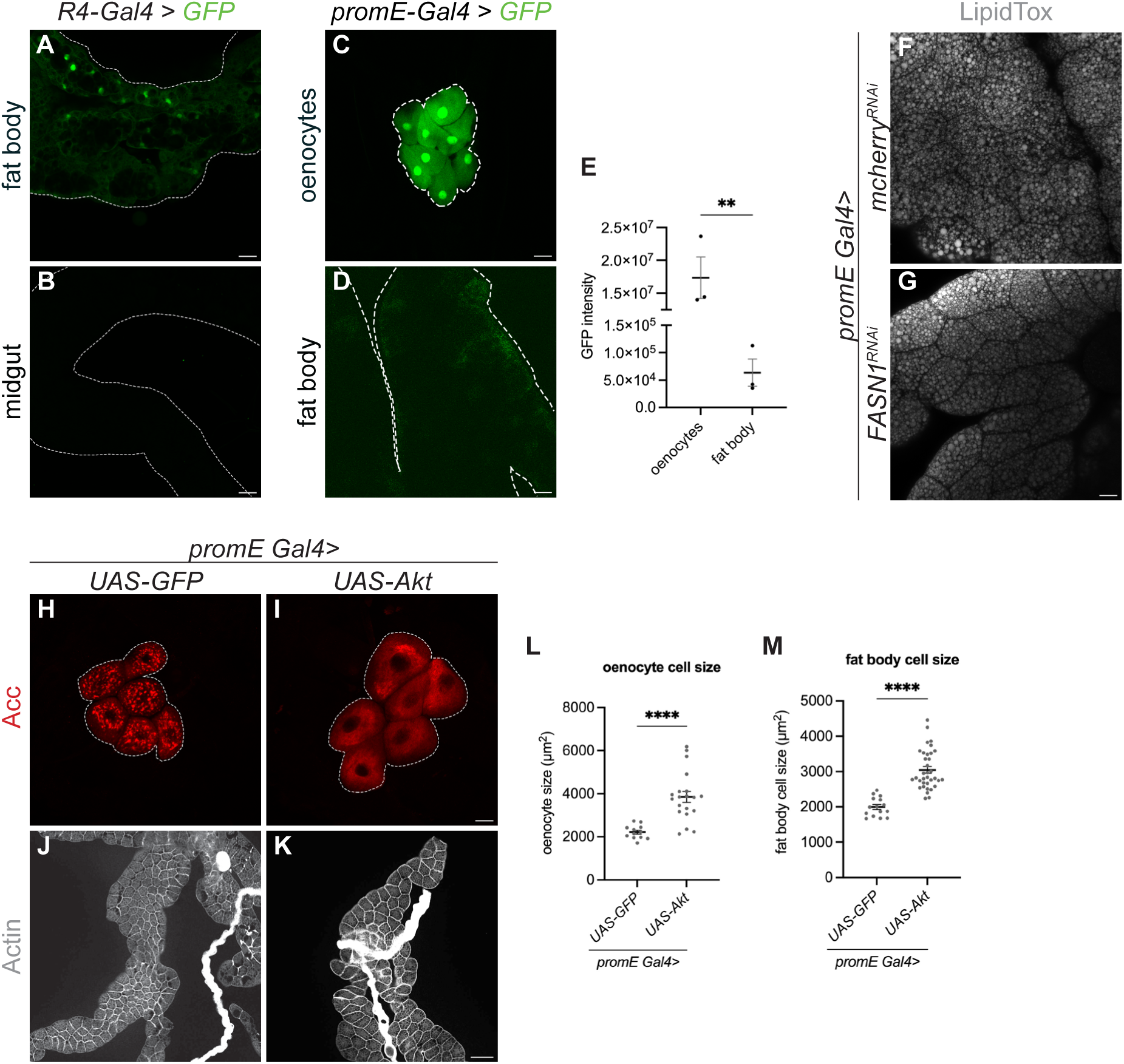
Specificity of R4 and promE Gal4, and oenocyte specific overexpression of PI3K signalling increase oenocytes and fat body cell size in healthy animals. **(A-B)** Representative image of the fat body (A, dashed lines) and midgut (B, dashed lines) from animals with *UAS-GFP* (green) driven by *R4-Gal4*. **(C-D)** Representative image of the oenocytes (A, dashed lines) and fat body (B, dashed lines) from animals with *UAS-GFP* (green) driven by *promE-Gal4*. **(E)** Quantification of GFP intensity in (A-B). Two-tailed unpaired t test: ** p = 0.0054. oenocytes: n = 3, mean ± SEM = 17367150 ± 3155724. fat body: n = 3, mean ± SEM = 63498 ± 24862. **(F-G)** Representative maximum projection of the LDs in fat body from healthy animals, where *mcherry^RNAi^* (F), *FASN1^RNAi^* (G) were expressed in the oenocytes. LipidTox (grey). **(H-I)** Representative maximum projection of the oenocytes (dashed lines) from healthy animals, where *UAS-GFP* (H) and *UAS-Akt* (I) were expressed in the oenocytes. Acc (red). **(J-K)** Representative maximum projection of the fat body cells from healthy animals, where *UAS-GFP* (J) and *UAS-Akt* (K) were expressed in the oenocytes. Fat body stained for phalloidin (Actin) (grey). **(L)** Quantification of oenocyte cell size in (H-I). Two-tailed unpaired t test: **** p < 0.0001. *UAS-GFP*: n = 12, mean ± SEM = 2215 ± 90.84. *UAS-Akt*: n = 20, mean ± SEM = 3855 ± 255.3. **(M)** Quantification of fat body cell size in (J-K). Two-tailed unpaired t test: **** p < 0.0001. *UAS-GFP*: n = 15, mean ± SEM = 1997 ± 69.20. *UAS-Akt*: n = 35, mean ± SEM = 3048 ± 94.12. Scale bar is 25μm in (C-D, F-I), 50 μm in (A-B), 100 μm in (J-K).

**Supplementary Figure 2.**
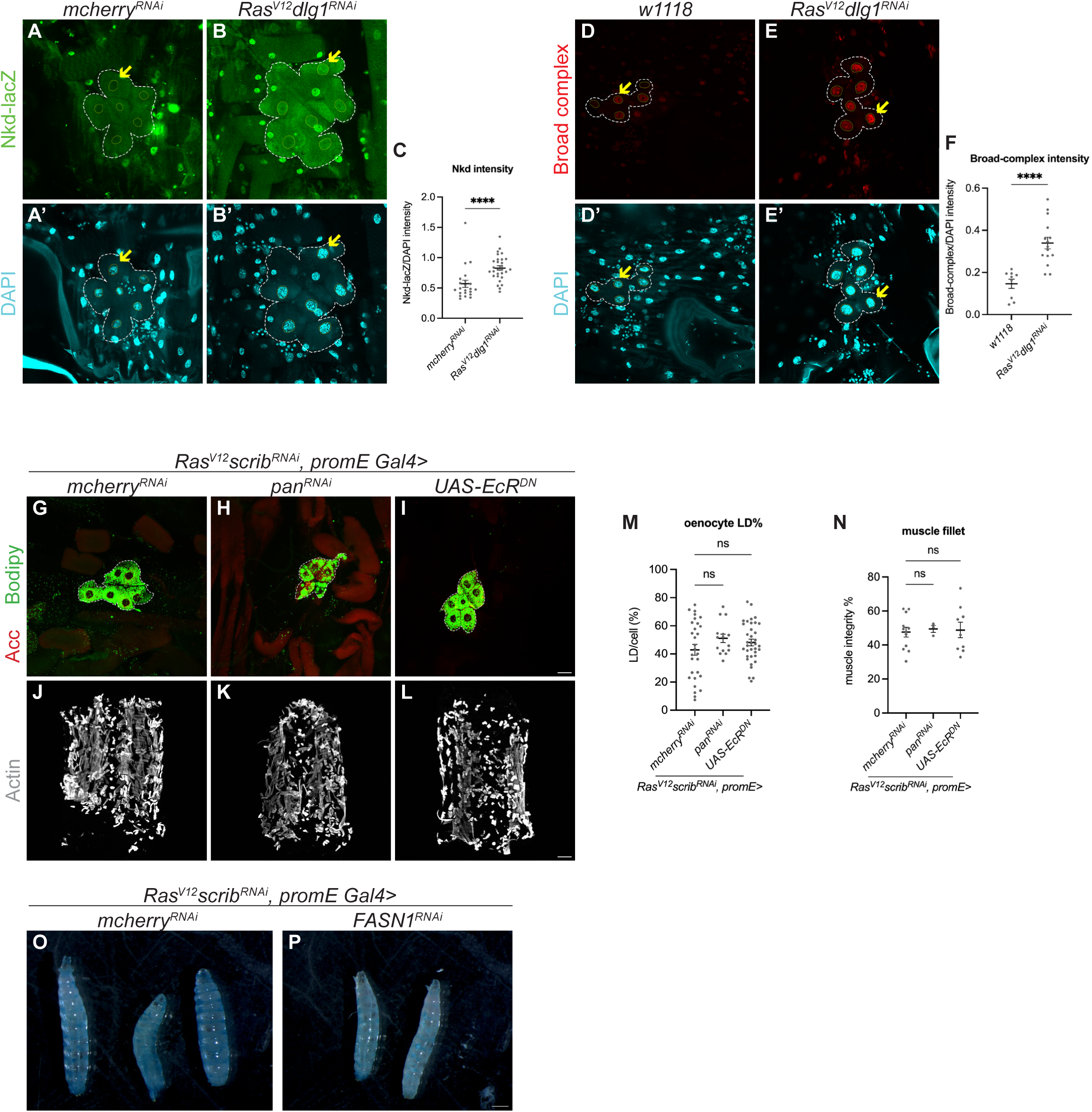
Wnt/wingless and ecdysone signalling pathways are altered in the oenocytes of cachectic animals. **(A-B’)** Representative maximum projections of the oenocytes (dashed lines) from *mcherry^RNAi^* (A-A’) and *Ras^V12^ dlg1^RNAi^*tumour-bearing animals (B-B’). Oenocytes stained for nkd-lacZ (green) in the nucleus (yellow arrows) (A-B), counterstained with DAPI (cyan) (A’-B’). **(C)** Quantification of oenocyte nkd-lacZ intensity normalised to DAPI intensity in (A-B’). Mann-Whitney test: **** p < 0.0001. *mcherry^RNAi^*: n = 22, mean ± SEM = 0.5704 ± 0.05908. *Ras^V12^ dlg1^RNAi^*: n = 29, mean ± SEM = 0.8288 ± 0.03864. **(D-E’)** Representative maximum projections of the oenocytes (dashed lines) from *w1118* (D-D’) and *Ras^V12^ dlg1^RNAi^* tumour-bearing animals (E-E’). Oenocytes stained for broad-complex (red) in the nucleus (yellow arrows) (D-E), counterstained with DAPI (cyan) (D’-E’). **(F)** Quantification of oenocyte broad-complex intensity normalised to DAPI intensity in (D-E’). Two-tailed unpaired t test: **** p < 0.0001. *w1118*: n = 9, mean ± SEM = 0.1464 ± 0.02195. *Ras^V12^ dlg1^RNAi^*: n = 15, mean ± SEM = 0.3397 ± 0.02769. **(G-I)** Representative maximum projections of the oenocytes (dashed lines) from *Ras^V12^ scrib^RNAi^* tumour-bearing animals, where *mcherry^RNAi^*(G), *pan^RNAi^* (H), *UAS-EcR^DN^* (I) were expressed in the oenocytes. Acc (red), Bodipy (green). **(J-L)** Representative images of the muscle fillet from *Ras^V12^ scrib^RNAi^* tumour-bearing animals, where *mcherry^RNAi^* (J), *pan^RNAi^* (K), *UAS-EcR^DN^* (L) were expressed in the oenocytes. Actin (grey). **(M)** Quantification of LD area as a percentage of individual oenocyte cell area in (G-I). Kruskal-Wallis test and Dunn’s test to correct for multiple comparisons: ns p = 0.4927. *mcherry^RNAi^*: n = 29, mean ± SEM = 42.84 ± 3.922. *pan^RNAi^*: n = 15, mean ± SEM = 51.03 ± 2.959. *UAS-EcR^DN^*: n = 36, mean ± SEM = 48.10 ± 2.354. **(N)** Quantification of muscle detachment in (J-L). Kruskal-Wallis test and Dunn’s test to correct for multiple comparisons: ns p = 0.9470. *mcherry^RNAi^*: n = 12, mean ± SEM = 47.54 ± 2.922. *pan^RNAi^*: n = 3, mean ± SEM = 49.47 ± 2.124. *UAS-EcR^DN^*: n = 9, mean ± SEM = 48.83 ± 4.584. (**O-P**) Trachea flooding assaying showing no blue liquid is incorporated in the trachea of *Ras^V12^ scrib^RNAi^* tumour bearing animals expressing *mcherry^RNAi^* (O) or *FASN1^RNAi^* (P) in the oenocytes. Scale bar is 25μm for oenocytes, 250μm for muscle fillet, 50mm for larvae in (O-P).

